# Quantitative dSTORM super-resolution microscopy localizes Aurora kinase A/AURKA in the mitochondrial matrix

**DOI:** 10.1101/2020.11.20.390948

**Authors:** Béatrice Durel, Charles Kervrann, Giulia Bertolin

## Abstract

Mitochondria are dynamic organelles playing essential metabolic and signaling functions in cells. Their ultrastructure has largely been investigated with electron microscopy (EM) techniques, which provided a wide range of information on how mitochondria acquire a tissue-specific shape, how they change during development, and how they are altered in disease conditions. However, quantifying protein-protein proximities using EM is extremely challenging. Super-resolution microscopy techniques as direct stochastic optical reconstruction microscopy (dSTORM) now provide a fluorescent-based alternative to EM with a higher quantitative throughput. Recently, super-resolution microscopy approaches including dSTORM led to valuable advances in our knowledge of mitochondrial ultrastructure, and in linking it with new insights in organelle functions. Nevertheless, dSTORM is currently used to image integral mitochondrial proteins only, and there is little or no information on proteins transiently present at this compartment. The cancer-related Aurora kinase A/AURKA is a protein localized at various subcellular locations, including mitochondria. After performing dSTORM, we here use the Geo-coPositioning System (GcoPS) image analysis method to quantify the degree of colocalization of AURKA with compartment-specific mitochondrial markers. We show that two-color dSTORM provides sufficient spatial resolution to visualize AURKA in the mitochondrial matrix. We conclude by demonstrating that optimizing fixation procedures is a key step to follow AURKA in the matrix. In this light, we show that a methanol-based fixation leads to a better detection of the matrix pool of AURKA than an aldehyde-based fixation. Our results indicate that dSTORM coupled to GcoPS colocalization analysis is a suitable approach to explore the compartmentalization of non-integral mitochondrial proteins as AURKA, in a qualitative and quantitative manner. This method also opens up the possibility of analyzing the proximity between AURKA and its multiple mitochondrial partners with exquisite spatial resolution, thereby allowing novel insights into the mitochondrial functions controlled by AURKA.

## Introduction

Mitochondria are multifunctional organelles involved in a wide range of cellular functions and signaling pathways. They are organized in an interconnected network spread throughout the cell surface, and in contact with several other subcellular compartments (van der Laan et al., 2016). Mitochondria are composed of two membranes: the Outer Mitochondrial Membrane (OMM), which encloses the organelle, and the Inner Mitochondrial Membrane (IMM), which is folded in invaginations called *cristae* and contains the oxidative phosphorylation system (OXPHOS). Two soluble compartments are also present: the Intermembrane Space (IMS), which separates the OMM from the IMM, and the matrix, which is the innermost compartment and where mtDNA molecules are located (Nunnari and Suomalainen, 2012).

In the past decades, our understanding of mitochondrial sub-architecture greatly benefited from advances in electron microscopy (EM). This technique turned out to be particularly useful in revealing how the two membranes are organized in various cell types and in tissues (reviewed in (Frey and Mannella, 2000)), and in highlighting how mitochondria are structurally altered in disease (Vincent et al., 2016) (Siegmund et al., 2018). However, EM is less convenient to explore the intramitochondrial localization of individual proteins. To be detected, antibodies specific to the protein of interest need to be coupled to gold beads – an approach known as immunogold EM –. Although this allows to visualize the sub-mitochondrial distribution of a protein with exquisite spatial resolution (Vogel et al., 2006), immunogold EM is a procedure with a relatively low efficiency. In individual EM sections, gold particles are generally scarce, and it is therefore nearly impossible to conclude on the localization of a protein by looking at single mitochondria. To extract quantitative information, individual localizations must be calculated from a considerable number of images before obtaining an average localization coefficient (Enger, 2017; Hayat, 1992). To understand protein associations in cells, double-immunogold EM is particularly convenient (Boykins et al., 2016). This consists of imaging two proteins within the same sample using gold beads of different sizes. With such approach, beads of 10 and 20 nm were used to detect the colocalization of mitochondrial complex III and IV in individual mitochondria (Golic et al., 2016). However, images of mitochondria with two sizes of gold beads are generally hard to interpret and to quantify, especially when beads are adjacent or clustered together.

Recent advances in fluorescence microscopy now offer elegant alternatives to electron microscopy. The width of a mitochondrion is between 200 and 500 nm, and the two mitochondrial membranes are 10-20 nm apart (Kaasik et al., 2007). Therefore, these organelles are beyond the resolution limit of most optical microscopes. This limit makes conventional fluorescence microscopy not suitable to explore the distribution of proteins within specific submitochondrial compartments. The recent development of super-resolution techniques allows to go beyond this diffraction limit, and it is now easier to investigate mitochondrial protein distribution in a quantitative manner (Jakobs and Wurm, 2014). Among super-resolution approaches, direct stochastic optical reconstruction microscopy (dSTORM) uses the transitions between the on- and off-states of fluorophores for multiple imaging cycles, which result in the activation/deactivation of fluorophores in a stochastic manner (Heilemann et al., 2008), (reviewed in (Samanta et al., 2019a)). A high-resolution map of individual fluorophores is obtained by recording the activation/deactivation rates of each fluorophore and their respective *xyz* coordinates. Given that this method provides an optical resolution of ∼10 nm, well beyond the diffraction limit, dSTORM immediately showed its potential for mitochondrial imaging. It was used to image mitochondria-microtubule contacts (Huang et al., 2008), the distribution of specific proteins as the ATP-synthase F1 α subunit or the MICOS complex (Dlasková et al., 2018; Stephan et al., 2020), and to explore the localization of mitochondrial nucleoids (Dlasková et al., 2018).

Integral mitochondrial proteins such as MICOS subunits showed very little or no extra-mitochondrial signal in dSTORM analyses (Stephan et al., 2020). As the non-mitochondrial signal comes from fluorophores not directly linked to the protein of interest, it turns on and off with different rates than mitochondria-bound fluorophores. Therefore, the extra-mitochondrial signal can be easily interpreted and treated as background noise when the protein is localized exclusively at mitochondria. However, discriminating between mitochondrial and extra-mitochondrial signal is a greater challenge when a given protein has both a mitochondrial and an extra-mitochondrial localization. This is the case of the Ser/Thr kinase AURKA, which has multiple subcellular locations including the centrosome and the mitotic spindle (reviewed in (Nikonova et al., 2013)), the nucleus (Zheng et al., 2016), and mitochondria (Bertolin et al., 2018; Grant et al., 2018). Using biochemical approaches, electron microscopy or molecular modeling, we and others provided evidence that AURKA localizes at mitochondria (Bertolin et al., 2018; Grant et al., 2018), and it is imported in the matrix (Bertolin et al., 2018). After import, AURKA was shown to regulate organelle dynamics and ATP levels throughout the cell cycle (Bertolin et al., 2018; Grant et al., 2018; Kashatus et al., 2011). Although it is conceivable that the regulation of these functions requires the interaction of AURKA with a great number of mitochondrial partners, we still have a fragmented view of the mitochondrial signaling cascade(s) in which AURKA is involved in. Understanding protein/protein proximities between AURKA and its interactors with a submitochondrial spatial and temporal resolution would constitute a significant step forward in understanding how AURKA orchestrates its multiple functions at this compartment.

We here show that the submitochondrial distribution of AURKA can be monitored with exquisite spatial resolution using two-color dSTORM. First, we evaluate the performance of two-color dSTORM in resolving OMM and IMM integral proteins within the same cell. We then optimize sample preparation procedures to visualize the presence of AURKA at mitochondria. Last, we exploit the quantitative potential of dSTORM and of rapid, noise-insensitive image processing methods to assess the colocalization of AURKA with OMM and matrix markers (Lavancier et al., 2019). Unlike electron microscopy, we show that this strategy allows to extract quantitative information on the intramitochondrial localization of AURKA directly from individual cells.

## Results and discussion

### Separating OMM and IMM using integral mitochondrial markers and quantitative dSTORM analyses

We first aimed at exploring whether dSTORM was a suitable approach to spatially distinguish the OMM from the IMM in MCF7 cells. To this end, we selected integral proteins with a known OMM or IMM localization, thereby allowing us to distinguish the two compartments unambiguously. We first fixed the cells with a mixture of paraformaldehyde/glutaraldehyde (PFA/G) to maintain mitochondrial morphology. We then labeled the OMM with an anti-TOMM22 primary antibody and we used it in combination with a primary antibody targeting the IMM marker COX2, or another targeting the IMM/matrix marker PMPCB. The anti-TOMM22 antibody was then immunodecorated with a secondary antibody conjugated to Alexa647, while anti-COX2 or anti-PMPCB antibodies were labeled with a secondary antibody conjugated to Alexa555. Two-color dSTORM recordings in 2D successfully managed to image TOMM22 together with COX2 (Fig. 1A), and with PMPCB (Fig. 1C). The TOMM22-specific staining appeared as a non-contiguous, ring-like structure enclosing individual mitochondria. This spatial distribution corroborates previous data obtained with Stimulated Emission Depletion (STED) nanoscopy in mammalian cell lines. In these reports, analyses performed in mammalian cells showed that the cognate TOMM20 protein organizes in clusters with variable density, and the size of these clusters can change according to the metabolic conditions of the cells (Wurm et al., 2011). When looking at internal submitochondrial compartments, COX2 and PMPCB were distributed as expected (Fig. 1A-C). However, both markers showed a diffused staining, and *cristae* were hardly visible. This could be due to sample preparation procedures, as the use of mild detergents facilitates the access of the structure of interest to antibodies on one hand, but it also destabilizes the lipid layer of the IMM on the other hand (Jacquemet et al., 2020). This could be a significant caveat when imaging the IMM using dSTORM. Whenever the integrity of the *cristae* is a mandatory parameter, high resolution live imaging approaches as STED could be envisaged (Stephan et al., 2019). However, it is important to remember that the distance between OMM and IMM normally falls between 15 and 25 nm, according to the model used (Reichert and Neupert, 2002). Since STED has an axial resolution less important than dSTORM (60 nm vs 20 nm, respectively (Samanta et al., 2019b)), employing STED microscopy could prevent an efficient separation of the OMM-and the IMM-specific signals.

**Fig. 1.**
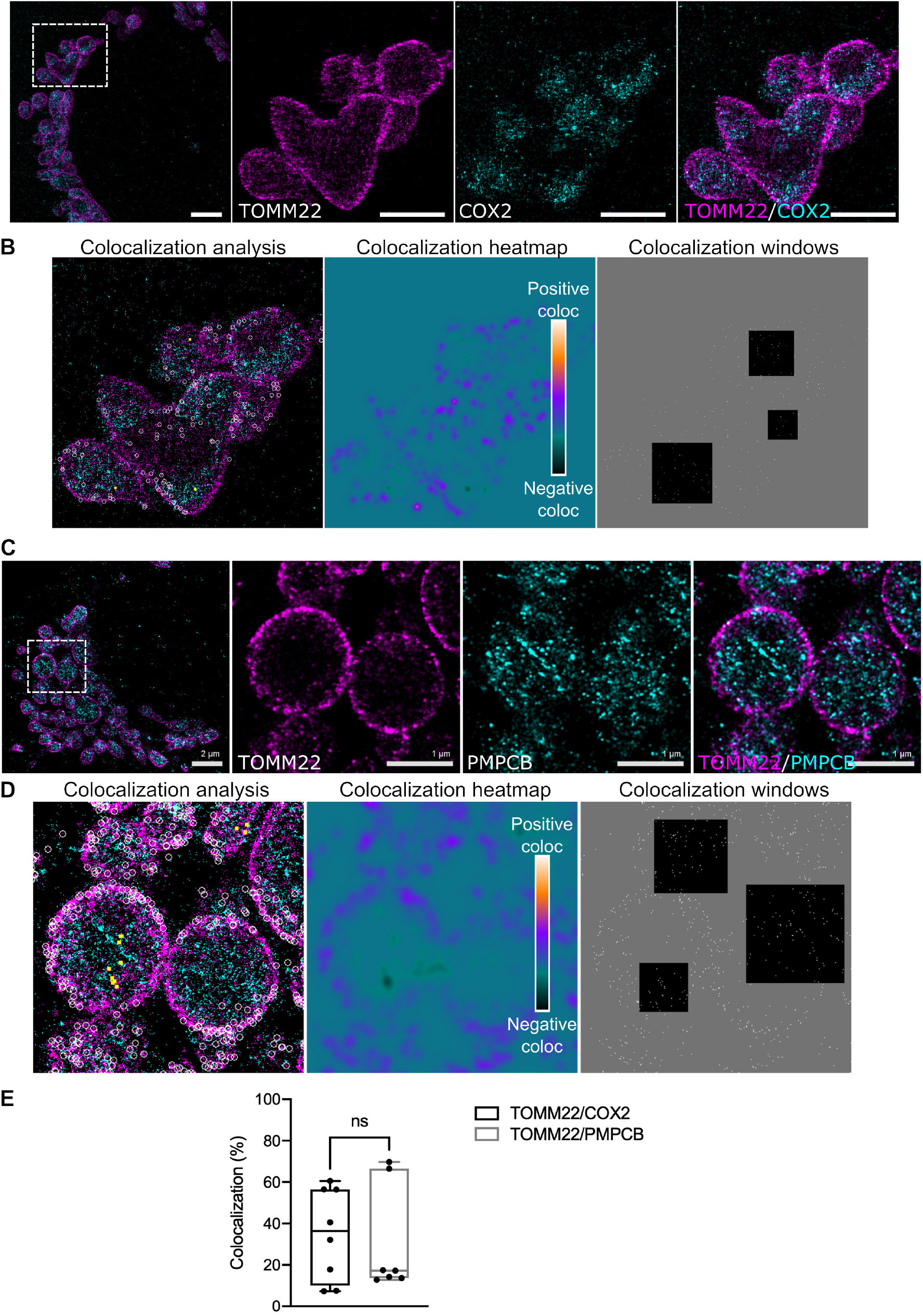
The OMM marker TOMM22 is separated from the IMM marker COX2 and the matrix protein PMPCB. **(A-C)** Maximal projections of representative 2D dSTORM micrographs from MCF7 co-stained for TOMM22 and COX2 (A), or for TOMM22 and PMPCB (C). The anti-TOMM22 primary antibody was detected using a secondary antibody conjugated to Alexa647 and pseudocolored magenta, while anti-COX2 and anti-PMPCB primary antibodies were detected using secondary antibodies conjugated to Alexa555 and pseudocolored cyan. The dotted area in the left panels indicates the magnified region where each staining is shown individually (middle panels), or merged (right panels). Scale bar: 2µm (left) or 1µm (middle and right panels). **(B-D)** (Left panels) Visual representation of positive (white rounds) and negative (yellow squares) colocalization sites on the magnified area of TOMM22/COX2 (B) and of TOMM22/PMPCB (D) micrographs after GcoPS analysis. (Middle panels) Colocalization heatmap ranging from yellow (positive colocalization) to black (negative colocalization), and background (colocalization score arbitrarily set to 0) in green. (Right panels). Three representative colocalization windows of 64 x 64 pixels, 96 x 96 pixels and 128 x 128 pixels superimposed over 3 representative positions. For each position, a window center is drawn in the set of super-localizations of the reference channel, and labeled as a white spot which is used for colocalization. All the window centers tested are shown. **(E)** Quantification of the percentage of colocalization for the indicated protein pairs. Data range from min to max. Dots correspond to individual cells issued from *n* = 3 independent experiments. ns = not significant

We then performed colocalization analyses to quantify the degree of colocalization between OMM and IMM/matrix markers. To this end, we used GcoPS, an object-based and intensity-based hybrid colocalization method which is rapid, insensitive to noise and capable of performing with comparable precision both on whole images and on image subregions (Lavancier et al., 2019). By assessing the degree of colocalization using this method, we estimate the likelihood of certain proteins to associate with specific mitochondrial sub-compartments at the nanoscale, and therefore deduce protein localization between the OMM and IMM. GcoPS takes as inputs two binary images computed from dSTORM super-localizations. In (Lavancier et al., 2019), the authors demonstrated that GcoPS provides reliable colocalization results and outperforms previous competitive methods like those of (Costes et al., 2004) and (Lagache et al., 2015) on images containing noise, irregularly shaped fluorescent patterns, and different optical resolutions. dSTORM micrographs are first translated into binary images with pixel size set to 10 nm. Then, colocalization is derived by approximating the interaction strength between the two proteins by the area of the intersection between the two binary images. A local-to-global approach is then used to quantify colocalization. First, the colocalization analysis is independently performed over squared windows randomly drawn in the whole image. The window size is uniformly drawn in the range from 64 x 64 to 128 x 128 pixels. Second, we derive p-values from multiple colocalization scores, which are further exploited to compute a global score from all the individual tests performed in the entire image. In our experiment, the commonly-used significance level alpha is set to 0.05 and the number of tested windows is set to 5000 for all the images analyzed. A lower or higher number of tests did not modify the statistical results, as shown in Table 1. From the sparse localizations of windows/tests/scores, we generated heatmaps to better visualize colocalizations on the entire image. For visualization purposes, the heatmaps were obtained by interpolating the score values using a Gaussian filter with a standard deviation of 50 nm. Colocalization analyses were performed in 2D, and this for two reasons: first, the lateral and axial dimensions of organelles in cell lines ensure the analysis of individual mitochondria within one observation volume and second, the signal/noise ratio is optimized on a larger number of particles than the few ones found on individual *z* slices.

At mitochondria, the colocalization coefficient for the TOMM22/COX2 and the TOMM22/PMPCB pairs is ∼30% (Fig. 1E). Globally, colocalization analysis obtained with the GcoPS method reveal that TOMM22 has a significant, although partial, degree of colocalization with two inner mitochondrial proteins. However, the overall colocalization coefficients between TOMM22 and the two IMM/matrix markers analyzed is similar, indicating that in our setup, dSTORM does not provide sufficient spatial resolution to separate the IMM from the matrix. However, within individual mitochondria, GcoPS detected areas where OMM and IMM/matrix proteins show anti-colocalization. These values correspond to the probability that two proteins are not in contact with each other (Fig. 1 B-D, left panels) (Bolte and Cordelieres, 2006). In addition, colocalization heatmaps show that the two protein pairs do not colocalize stochastically throughout the two mitochondrial membranes, but colocalization is rather concentrated in discrete proximity sites between the OMM and the IMM/matrix (Fig. 1 B-D, middle panels). These sites are reminiscent of mitochondrial membranes “contact sites” (Reichert and Neupert, 2002), although it should be kept in mind that the steric hindrance of primary and secondary antibodies used in dSTORM could alter the overall number, the diameter and volume of these proximity sites.

Our results show that dSTORM is a convenient approach to separate the OMM from the IMM/matrix compartments using well characterized, integral mitochondrial proteins. Therefore, this technique could be used to monitor the presence of non-integral mitochondrial proteins and their relative submitochondrial distribution.

### dSTORM identifies AURKA in the mitochondrial matrix of methanol-fixed cells

After establishing that dSTORM is a suitable approach to localize mitochondrial integral proteins, we sought to explore whether it is powerful enough to determine the submitochondrial localization of proteins with multiple cellular locations. This is a challenge, as the extra-mitochondrial signal, or the fraction of the protein undergoing import inside the organelles, could hide its ultimate submitochondrial location. The multifunctional kinase AURKA is an example of a protein localizing in multiple cellular compartments at a time. AURKA was shown to localize at mitochondria (Bertolin et al., 2018; Grant et al., 2018) and to interact with the TOMM machinery prior to import in the mitochondrial matrix (Bertolin et al., 2018). Its localization in the matrix was shown using immunogold beads and TEM after overexpressing AURKA in cells, which also allowed to detect a minor fraction of the protein at the OMM. However, we found the signal corresponding to endogenous AURKA to be weak and difficult to detect in our previous immunogold EM analyses (Bertolin et al., 2018), implying that TEM experiments can be performed only upon AURKA overexpression. Given that the overexpression of AURKA induces several major alterations to cell physiology (Nikonova et al., 2013; Bertolin and Tramier, 2019), we asked whether we could improve the detection of endogenous AURKA using alternative, fluorescence-based techniques. Therefore, we performed dSTORM analyses to assess the submitochondrial localization of AURKA and to test its proximity with key mitochondrial components.

To detect endogenous AURKA, we used the monoclonal anti-AURKA 35C1 clone. This antibody was previously shown to be efficient in immunofluorescence and biochemical approaches, and to measure the activity of the kinase *in vivo* (Cremet et al., 2003). With this tool, we explored the colocalization of AURKA with OMM and IMM markers. Due to the species cross-reactivity of the anti-AURKA 35C1 antibody, we could not use an anti-TOMM22 primary antibody as an OMM marker. Given that TOMM22 and TOMM20 both belong to the Translocase of Outer Membrane complex and interact with each other (van Wilpe et al., 1999), we turned to a compatible anti-TOMM20 antibody to label the OMM. We first confirmed the suitability of TOMM20 as a potential OMM reference for AURKA, by comparing the colocalization between TOMM22 and TOMM20 or another OMM protein, VDAC1. Anti-TOMM20 and anti-VDAC1 antibodies were detected with a secondary antibody conjugated to Alexa555, while the anti-TOMM22 antibody was conjugated using a secondary antibody conjugated to Alexa647 as in Fig. 1. Previous analyses obtained with STED microscopy revealed that VDAC1 is not uniformly distributed throughout the OMM, but it rather clusters in TOMM-positive distinct domains (Neumann et al., 2010). We retrieved a similar distribution of VDAC1 in clusters, and the colocalization degree with TOMM22 was indeed partial and did not exceed 25% (Fig. 2A-B, E). Therefore, our combined approach relying on dSTORM and GcoPS-based analyses confirmed previous results obtained with STED (Neumann et al., 2010). Conversely, the colocalization degree between the two TOMM subunits TOMM22 and TOMM20 was significantly increased, and it reached 60% for the majority of the cells analyzed (Fig. 2 C-D, E). This substantiates the choice of TOMM20 as an alternative TOMM subunit to be used for colocalization analyses with an anti-AURKA antibody.

**Fig. 2.**
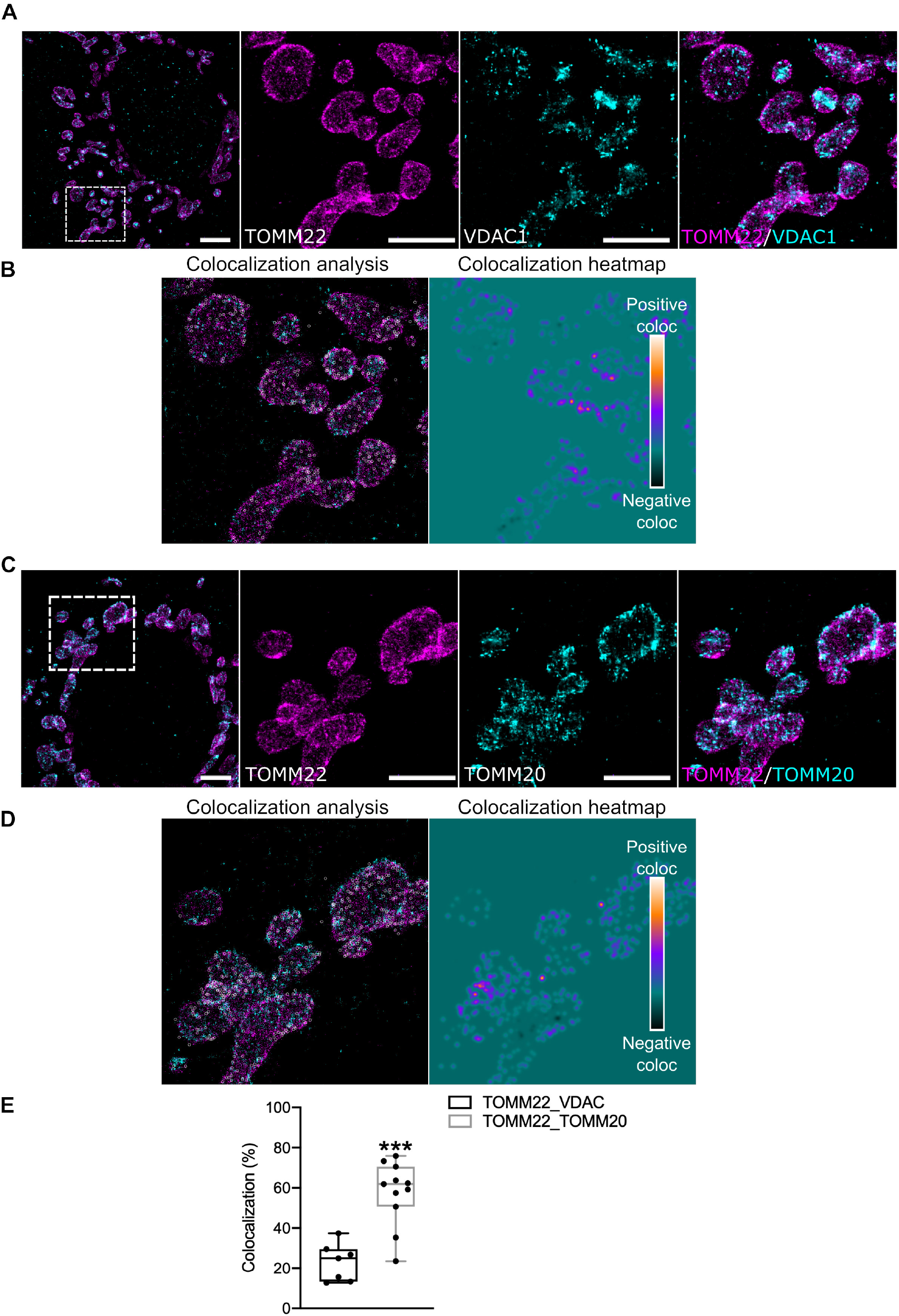
The OMM marker TOMM22 preferentially colocalizes with TOMM20 on the OMM. **(A-C)** Maximal projections of representative 2D dSTORM micrographs from MCF7 co-stained for TOMM22 and VDAC1 (A), or for TOMM22 and TOMM20 (C). The anti-TOMM22 primary antibody was detected using a secondary antibody conjugated to Alexa647 and pseudocolored magenta, while anti-VDAC1 and anti-TOMM20 primary antibodies were detected using secondary antibodies conjugated to Alexa555 and pseudocolored cyan. The dotted area in the left panels indicates the magnified region where each staining is shown individually (middle panels), or merged (right panels). Scale bar: 2µm (left) or 1µm (middle and right panels). **(B-D)** (Left panels) Visual representation of positive (white rounds) colocalization sites on the magnified area of TOMM22/VDAC1 (B) and of TOMM22/TOMM20 (D) micrographs after GcoPS analysis. (Right panels) Colocalization heatmap ranging from yellow (positive colocalization) to black (negative colocalization), and background (colocalization score arbitrarily set to 0) in green. **(E)** Quantification of the percentage of colocalization for the indicated protein pairs. Data range from min to max. Dots correspond to individual cells issued from *n* = 3 independent experiments. *** *P* < 0.001

In addition to TOMM20 and as shown in Fig. 1, PMPCB was used to label the mitochondrial matrix. Anti-TOMM20 and anti-PMPCB antibodies were detected using a secondary antibody conjugated to Alexa647, while the anti-AURKA 35C1 antibody was detected with a secondary antibody conjugated to Alexa555 (Fig. 3A, C). GcoPS-based analyses revealed that the colocalization between AURKA and TOMM20 does not exceed 17% (Fig. 3B, E), while the one between AURKA and PMPCB is 2-fold higher and nearly reaches 38% (Fig. 3D, E). The AURKA/PMPCB colocalization coefficients are in line with our previous observations showing that approximately 35-40% of endogenous AURKA is found as a processed form at mitochondria (Bertolin et al., 2018). These data also confirm our previous TEM analyses made with exogenous AURKA, thereby localizing the kinase mainly in the mitochondrial matrix. In addition, our results suggest that colocalization values obtained with GcoPS can be directly compared to molecular fractions obtained with biochemical approaches. Last, they open up the possibility of exploring the colocalization between endogenous AURKA and its potential substrates at mitochondria, a paradigm closer to cell physiology.

**Fig. 3.**
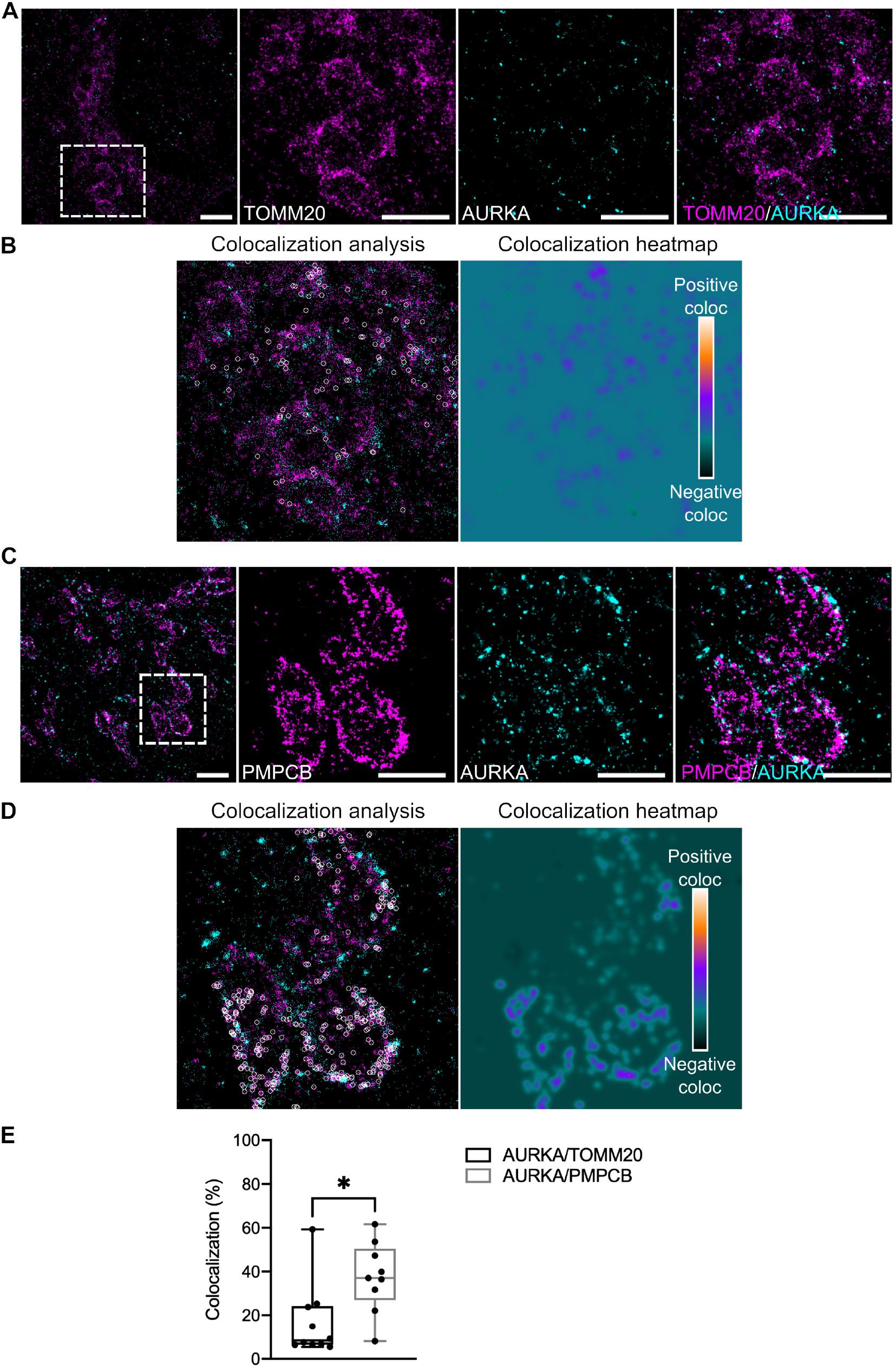
AURKA colocalizes with PMPCB in methanol-fixed cells. **(A-C)** Maximal projections of representative 2D dSTORM micrographs from MCF7 co-stained for AURKA and TOMM20 (A), or for AURKA and PMPCB (C). The anti-TOMM20 and anti-PMPCB primary antibodies were detected using secondary antibodies conjugated to Alexa647 and pseudocolored magenta, while the anti-AURKA primary antibody was detected using a secondary antibody conjugated to Alexa555 and pseudocolored cyan. For both protein pairs, the dotted area in the left panels indicates the magnified region where each staining is shown individually (middle panels), or merged (right panels). Scale bar: 2µm (left) or 1µm (middle and right panels). **(B-D)** (Left panels) Visual representation of positive (white rounds) colocalization sites on the magnified area of AURKA/TOMM20 (B) and of AURKA/PMPCB (D) micrographs. (Rght panels) Colocalization heatmap ranging from yellow (positive colocalization) to black (negative colocalization), and background (colocalization score arbitrarily set to 0) in green. **(E)** Quantification of the percentage of colocalization for the indicated protein pairs. Data range from min to max. Dots correspond to individual cells issued from *n* = 3 independent experiments. * *P* <0.05.

Our data indicate that dSTORM is a useful approach to follow the proximity of AURKA, a non-integral mitochondrial protein, with OMM and matrix components. Importantly, they separate two pools of AURKA at mitochondria, both qualitatively and quantitatively. While the larger pool of the kinase is located in the matrix, there is a smaller pool of AURKA which has an OMM localization and which gives a low, albeit positive, colocalization score. The two pools are not mutually exclusive, as AURKA needs to be imported through the OMM before reaching the mitochondrial innermost compartment (Bertolin et al., 2018). Altogether, our results underline that dSTORM coupled to GcoPS is capable of detecting small pools of the kinase with a potential functional readout.

### PFA/G fixation coupled to antigen retrieval procedures preserves mitochondrial ultrastructure, but it compromises the localization of AURKA in the matrix

As the anti-AURKA 35C1 primary antibody retrieves no AURKA-specific signal when cells are fixed using PFA/G, a methanol-based fixation is mandatory (Cremet et al., 2003). Despite the capacity of dSTORM to evaluate the proximity of AURKA with mitochondrial proteins in methanol-fixed cells, we reasoned that methanol might induce a partial loss of mitochondrial protein content, due to its lipid-dissolving action (Hoetelmans et al., 2001; Jamur and Oliver, 2010). Therefore, we evaluated the performance of methanol-and PAF/G-based fixation methods. When comparing the effect of the two fixation procedures on TOMM20 and PMPCB, we noticed a loss in the quality of both stainings. The distribution of TOMM20 was similar to the one of TOMM22 in PFA/G-fixed cells, with a well-defined ring-like structure and a non-contiguous staining (Fig. 4A, compare with Fig. 1A). On the contrary, the OMM appeared ruptured, and with an irregular TOMM20 staining in methanol-fixed cells. A similar loss in mitochondrial content was also observed for PMPCB, which appeared fragmented and not contiguous upon fixation with methanol (Fig. 4B). On the contrary, the matrix appeared intact when cells were fixed with PFA/G (Fig. 4B). As shown above, dSTORM coupled to GcoPS was able to retrieve a large pool of AURKA colocalizing with the matrix marker PMPCB. However, it should be kept in mind that harsh fixation methods as methanol may underestimate or even prevent the visualization of protein proximities. This is due to the partial loss in mitochondrial protein content that is induced by methanol. On one hand, TOMM20 and PMPCB are abundant mitochondrial proteins and colocalization coefficients can still be calculated on the pool of the proteins preserved after fixation. On the other hand, it could be challenging – or even impossible – to evaluate the proximity between AURKA and mitochondrial partners which have lower expression levels. Therefore, we reasoned that improvements in sample preparation procedures are required to preserve mitochondrial ultrastructure while visualizing the kinase and its potential interactors at these organelles.

**Fig. 4.**
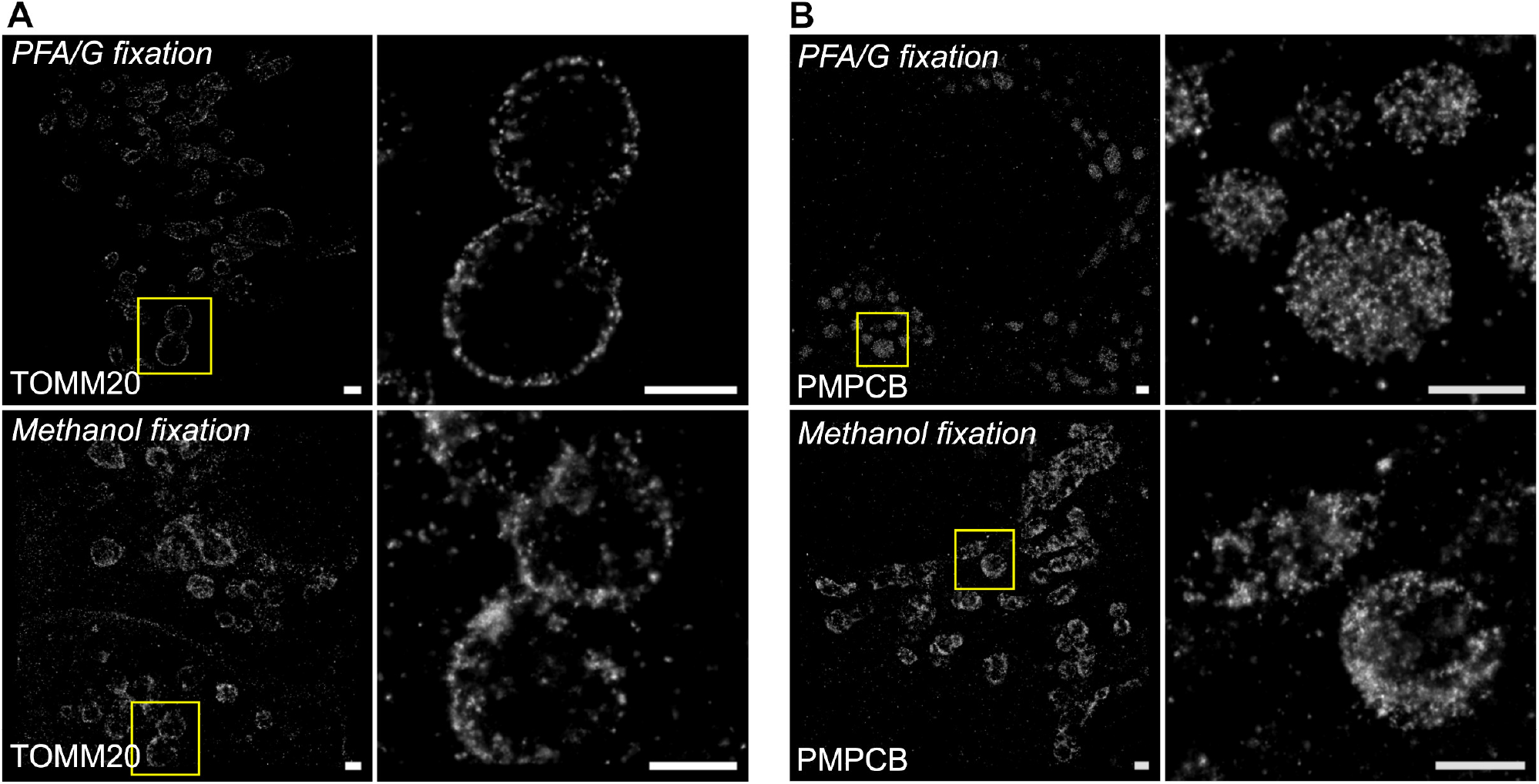
Methanol fixation degrades mitochondrial ultrastructure. (A-B) Representative images of MCF7 cells stained for TOMM20 (A) or PMPCB (B) after being fixed in a PFA/G mixture (upper panels) or in ice-cold methanol (lower panels). Primary antibodies were detected using secondary antibodies conjugated to Alexa647. The yellow insets indicate the magnified region for each protein. Scale bar: 1µm.

To improve the visualization of AURKA at mitochondria, we reasoned that our staining protocol needed to be improved on two aspects. First, we needed a fixation method to preserve organellar ultrastructure. Then, antigen retrieval methods were required to expose the epitope of AURKA recognized by the 35C1 primary antibody, which is normally hidden by PFA/G. This procedure would allow us to detect the kinase at mitochondria, while using a fixation method able to maintain the integrity of mitochondrial membranes and proteins. In light of these considerations, we fixed MCF7 cells in PFA/G, as previously done for the TOMM22/COX2 and the TOMM22/PMPCB pairs. We then performed antigen retrieval procedures by incubating cells in citrate buffer for 15 min at 98°C, and this before proceeding with sample preparation for dSTORM. As in methanol-fixed cells, we combined the anti-AURKA 35C1 primary antibody with primary antibodies labelling the OMM marker TOMM20, or the matrix marker PMPCB. The anti-AURKA antibody was then detected with an Alexa555-conjugated secondary antibody, while the anti-TOMM20 or anti-PMPCB antibodies were detected with a Alexa647-conjugated secondary antibody. By performing dSTORM microscopy, our first observation was that the anti-AURKA 35C1 primary antibody detected mitochondria-like structures in MCF7 cells (Fig. 5A, C). Indeed, these structures could be juxtaposed with either the TOMM20 or the PMPCB staining, strongly indicating that the AURKA-positive signal detected under these fixation conditions corresponds to mitochondria. We then ran the GcoPS procedure to determine the degree of colocalization between AURKA and each of the two mitochondrial markers. GcoPS-based analyses revealed that the colocalization coefficients of the AURKA/TOMM20 and of the AURKA/PMPCB pairs are similar (Fig. 5B, D-E). Therefore, our analyses in cells subjected to PFA/G fixation and antigen retrieval procedures did not provide additional evidence in support of the matrix localization of AURKA. Although this fixation procedure globally ameliorated mitochondrial ultrastructure, it did not refine the colocalization data obtained in methanol-fixed cells, nor previous biochemical and electron microscopy data (Bertolin et al., 2018). A potential explanation for these results could be that PFA/G coupled to antigen retrieval procedures restored only a fraction of the antigens detected by the anti-AURKA 35C1 primary antibody. Most likely, the antigens located on the OMM are the most accessible ones and were retrieved first, while the antigens located in the matrix were more difficult to gain access to. This hypothesis is corroborated by the observation that colocalization coefficients for the AURKA/TOMM20 pair were ∼16% both in methanol-and PFA/G-fixed cells and were consistent with previous TEM data, while the AURKA/PMPCB colocalization coefficients were significantly lowered in PFA/G-fixed cells (14.3%) compared to those obtained in methanol-fixed cells (37.5%) and TEM analyses.

**Fig. 5.**
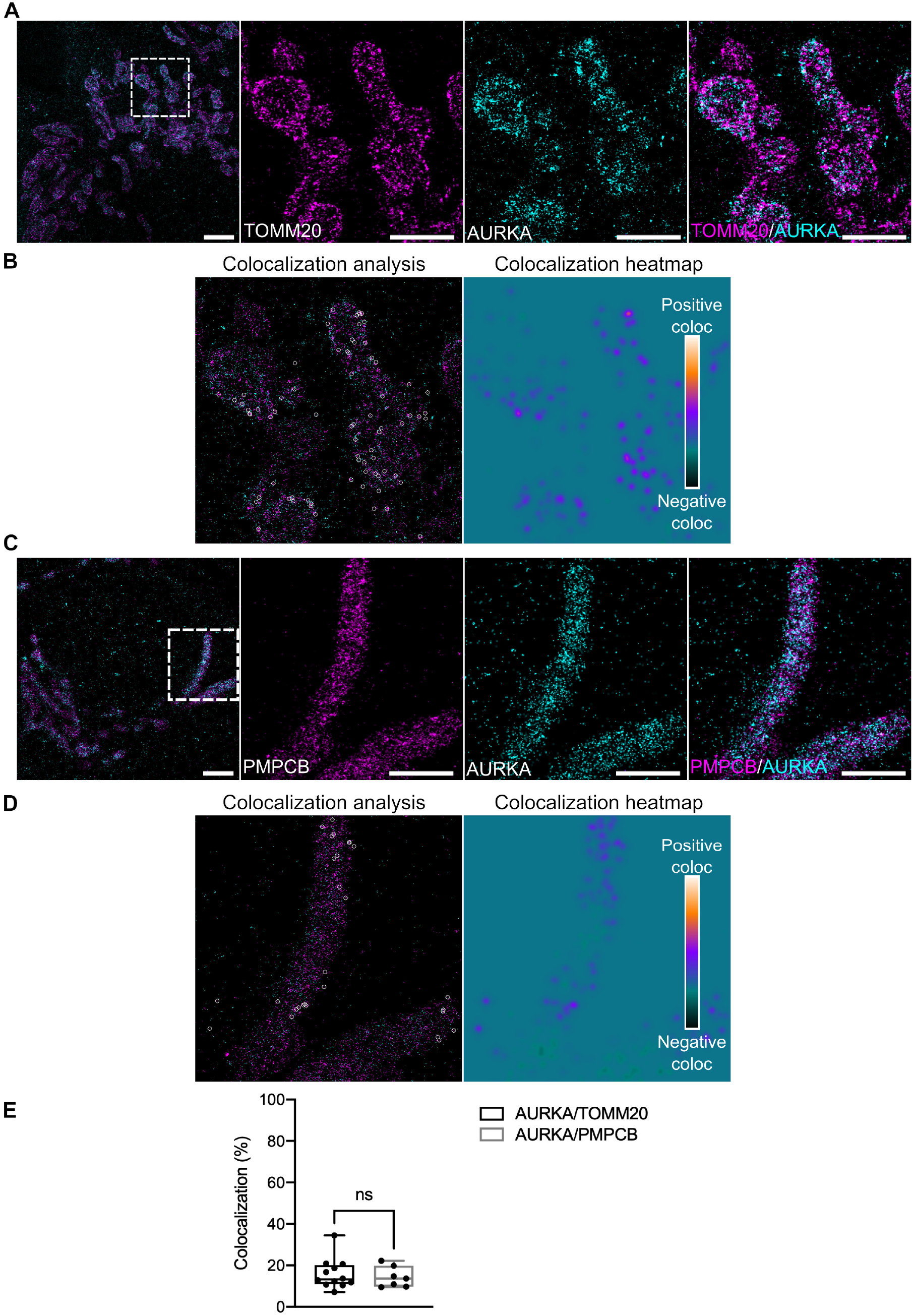
The preferential localization of AURKA in the matrix is not retrieved upon PFA/G fixation. **(A-C)** Maximal projections of representative 2D dSTORM micrographs from MCF7 co-stained for AURKA and TOMM20 (A), or for AURKA and PMPCB (B). The anti-TOMM20 and anti-PMPCB primary antibodies were detected using secondary antibodies conjugated to Alexa647 and pseudocolored magenta, while the anti-AURKA primary antibody was detected using a secondary antibody conjugated to Alexa555 and pseudocolored cyan. For both protein pairs, the dotted area in the left panels indicates the magnified region where each staining is shown individually (middle panels), or merged (right panels). Scale bar: 2µm (left) or 1µm (middle and right panels). **(B-D)** (Left panels) Visual representation of positive (white rounds) colocalization sites on the magnified area of AURKA/TOMM20 (B) and of AURKA/PMPCB (D) micrographs. (Right panels) Colocalization heatmap ranging from yellow (positive colocalization) to black (negative colocalization) and background (colocalization score arbitrarily set to 0) in green. **(E)** Quantification of the percentage of colocalization for the indicated protein pairs. Data range from min to max. Dots correspond to individual cells issued from *n* = 3 independent experiments. ns = not significant.

Overall, our data identify dSTORM as a qualitative and quantitative approach to investigate the localization of AURKA at mitochondria with a sub-organellar resolution. Using AURKA as an example of proteins with multiple subcellular locations, our results also shed light on how to sample preparation procedures are key to assess the vicinity of AURKA with specific mitochondrial proteins, and explore the potential consequences of alternative fixation methods on protein colocalization at mitochondria.

## Conclusions

Our data show that dSTORM is a highly appropriate approach to separate the different mitochondrial subcompartments with sufficient spatial resolution, and to follow the submitochondrial location of integral mitochondrial proteins. This confirms a recent elegant report, where two-color dSTORM and MINFLUX were used to follow the distribution of MICOS subunits throughout mitochondrial cristae (Stephan et al., 2020).

To the extent of our knowledge, we also show for the first time that dSTORM is suitable to detect non-integral mitochondrial proteins as AURKA, which have multiple subcellular locations and are only transiently present at this compartment. This indeed confirms that dSTORM provides an exquisite spatial resolution, but it also indicates that sample preparation procedures must be carefully optimized to preserve subcellular structures while detecting the protein(s) of interest. Despite these great advantages, it should also be noted that the temporal resolution of dSTORM is limited, mostly due to the composition of blinking buffers and imaging conditions which are largely incompatible with living samples. In this light, the recent development of MINFLUX appears to be a very promising alternative, combining a high spatial resolution with the possibility of using living samples (Balzarotti et al., 2017).

In terms of investigating mitochondrial functions, dSTORM is a useful method to define the submitochondrial locations of proteins with unprecedented resolution. The MitoCarta repository – the largest public database annotating mitochondrial proteins (Calvo et al., 2016) – was recently found to contain only half of the mitochondrial proteome (Antonicka et al., 2020). Bio-ID-based proteomics techniques significantly expanded our view of the mitochondrial proteome (Antonicka et al., 2020), including ER-mitochondrial contacts (Cho et al., 2020; Kwak et al., 2020). However, they lack in showing the distribution of proteins within the organelles, which could be helpful in inferring their function(s) at this compartment. In particular, this would represent a huge advantage for proteins that are still poorly characterized from the functional point of view. The orthogonal validation of selected hits – obtained with large-scale omics – with *in cellulo* dSTORM could represent a new frontier to infer on protein functions in specific submitochondrial compartments.

Last, the use of dSTORM provides new perspectives on the roles played by mitochondrial AURKA. Not only it allows to observe the localization of the kinase within single mitochondria, but it also allows to evaluate its physical proximity with new/potential substrates. This would undoubtedly provide researchers with an unprecedented insight on the multiple roles of AURKA at mitochondria. However, it should be kept in mind that the proximity of AURKA with potential substrates or interactors, should also be corroborated with complementary information on the activation status of the kinase. AURKA undergoes activation upon autophosphorylation on Thr288 (Bayliss et al., 2003; Cheetham, 2002; Zhang et al., 2007) before phosphorylating its substrates, both at mitosis and during interphase (reviewed in (Bertolin and Tramier, 2019; Nikonova et al., 2013)). We know that the kinase is active at mitochondria, thanks to a Förster’s Resonance Energy Transfer (FRET) biosensor monitoring activated AURKA in single cells (Bertolin et al., 2016). A fascinating future challenge could be to simultaneously monitor the activation of AURKA and its proximity with a putative substrate not in single cells, but in single mitochondria. Coupling FRET with dSTORM or, more generally, super-resolution microscopy could pave the way to the establishment of the mitochondrial “activome” of AURKA. In a therapeutic perspective, this functional network could be a promising tool for the treatment of patients with epithelial or hematological malignancies linked to the overexpression of AURKA.

## Materials and Methods

### Cell culture, fixation and antigen unmasking procedures

Mycoplasma-free MCF7 cells (HTB-22) were purchased from the American Type Culture Collection (ATCC) and grown in Dulbecco’s modified Eagle’s medium (DMEM, Life Technologies Thermo Fisher Scientific), supplemented with 10% Fetal Bovine Serum (FBS, Life Technologies Thermo Fisher Scientific) and 1% penicillin/streptomycin (Life Technologies Thermo Fisher Scientific) were cultured in 6-well plates on 1.5H 22 x 22mm coverslips (Marienfeld VWR ref: 630-2186), and then fixed in a mixture of methanol-free 4% paraformaldehyde (PFA, Euromedex EM-15710) and 0.2% Glutaraldehyde (Euromedex, EM-16221) in PBS 1X (Merck, P4417-100TAB), at 37°C for 20 min, or with ice-cold methanol at -20°C for 20 min. The autofluorescence generated by Glutaraldehyde was neutralized by using 1 mg/ml sodium borohydride at room temperature for 10 min (Sigma-Aldrich, 452882). AURKA antigen retrieval was performed by incubating cells previously fixed in 4% PFA/0.2% Glutaraldehyde in a 10 mM sodium citrate tribasic dihydrate solution (Sigma-Aldrich, C8532), pH 6 at 98°C on a hot plate for 15 min. To avoid evaporation, approximately 5 ml/well of this solution were used. After letting the plate cool down for 20 min at room temperature, the cells were washed twice in PBS 1X with gentle rocking, then permeabilized with 0.1% Triton X-100 (Sigma, 93 443-100ml) for 10 min. Cells were then saturated with 5% Bovine Serum Albumin (BSA) (Merck, A4503 50G) in PBS 1X for 30 min, with gentle rocking.

### Immunocytochemistry procedures

Primary antibodies were as follows: monoclonal mouse anti-AURKA (Clone 5C3; (Cremet et al., 2003) at 1:20 and anti-TOMM22 (Abcam, ab10436) at 1:5000; monoclonal rabbit anti-TOMM2O (Abcam, ab186734) at 1:500; polyclonal rabbit anti-PMPCB (Proteintech, 16064-1-AP) at 1:400, VDAC1 (Abcam, ab15895) at 1:1000 and used after antigen retrieval procedures, and COX2 (Agier et al., 2012) at 1:2000. All primary antibodies were diluted in 5% PBS/BSA and incubated overnight at 4° C. After 3 washes in 5% PBS/BSA for 10 min each with gentle rocking, AURKA, VDAC1 or COX2 were revealed by a species-specific Fab’ secondary antibody coupled to Alexa 555 (Thermo Fisher Scientific, A-21430 or A-21425). TOMM22 was revealed with a species-specific Fab’ secondary antibody coupled to Alexa 647 (Thermo Fisher Scientific, A-21237 or A-21246). TOMM20 and PMPCB were revealed with either secondary antibody depending on the co-stained protein. Both secondaries were used at a concentration of 1:1000 in 5% PBS/BSA and incubated in for 1 hour at 37° C.

The cells were then washed 3 times for 10 min in 1X PBS, then post-fixed in 4% PFA (Merck, 8187081000) at room temperature for 5 min, washed 3 times in 1X PBS and incubated for 5 min in 50mM ammonium chloride (Sigma-Aldrich, 254134) before mounting on cavity blades (Marienfeld, VWR, 630-1611), replenished with dSTORM buffer (Smart Kit, Abbelight) and sealed with dental paste (Rotec, Picodent twinsil speed 22, Ref: 13001002) to avoid oxygen exposure, immediately prior to observation.

### Image acquisition and reconstruction procedures

Two-color dSTORM acquisitions were performed on an Abbelight super resolution microscope (Abbelight, France) constituted by an Olympus IX83 microscope, an Abbelight SAFe 180 nanoscopy module, two Oxxius lasers at 640 nm and at 532 nm (Oxxius, France), a pco.panda sCMOS camera (PCO AG, Germany), equipped with a 100X oil-immersion objective (NA 1.5), and driven by the NEO software (Abbelight, France). Particle quantification and the reconstruction of the super resolution images was performed with the NEO software, using the Gaussian fitting algorithm integrated within. To evaluate the effect of the fixation method on mitochondrial integrity, TOMM20 or PMPCB-specific images were acquired with a home-made dSTORM microscope, composed of a Nikon TIRF system, a 640 nm laser (Toptica, Germany), an EMCCD camera (Andor, UK) and controlled by the NIS software. A 100X objective (NA 1.49) and a 1.5X lens were used to acquire images. Images were reconstructed using the UNLOC plugin for the ImageJ software (ImageJ, NHI), (Mailfert et al., 2018). 30000 frames of approximately 250 x 250 pixels were analyzed with the parameter-free UNLOC [UNsupervised particule LOCalization algorithm v1.0] algorithm in the high-density mode, allowing to fit with a multiple emitter fitting model, to correct the local and temporal variations of the background. The *xy* drift was corrected using a robust correlation method, and data were filtered according to these parameters (No filters parameters were modified except SNR Min = 20). Finally, an Integrated-Gaussian mode was applied to render the reconstructed images with a sub-pixel zoom factor of 8, leading to a final pixel size of 107 nm.

### Colocalization analyses and statistics

The colocalization method takes as inputs two binary images corresponding to dSTORM super-localizations translated into 0-1 images with a spatial resolution of 10 nm. The molecules with a localization uncertainty higher than 15 nm were discarded *a priori*. Colocalization was then performed on small squared windows as follows: the colocalization score is derived by approximating the interaction strength between the two proteins by the area of the intersection between the two binary images. In (Lavancier et al., 2019), this score is normalized with respect to the variance and is proved to follow a standard normal distribution. Therefore, the two proteins are expected to colocalize in the window of interest if the p-value is lower than the significance level α (α = 0.05). This procedure is then repeated *N* times by applying the colocalization test independently on square windows randomly drawn in the whole image. The size of patches is uniformly drawn in the ranges 64 x 64 to 128 x 128 pixels on dSTROM images and 21 x 21 to 42 x 42 pixels on Airycan images, respectively. Last, a global colocalization score is defined as the ratio of positive responses in windows over the total number *N* of tested windows. We considered *N* = 5000 tests for dSTORM images.

### Statistical analyses

After testing data for normality, the Mann-Whitney test (Fig. 1E, 2E, 3E and 5E) was used to compare GcoPS colocalization scores in each condition.

## Author Contributions

B.D. developed and optimized mitochondria-and AURKA-specific staining procedures, performed the experiments and particle quantification and reconstruction; C.K. optimized the GcoPS software for mitochondrial images, performed colocalization analyses and reviewed the manuscript; G.B. designed research, provided funding and wrote and revised the manuscript. All the authors provided feedback on the manuscript and agreed on its final version.

## Conflict of Interest

The authors declare no conflict of interest.

## Acknowledgments

We thank M. Tramier, X. Pinson and R. F. Laine for their helpful comments and feedback on the present manuscript. The authors thank the Abbelight company for providing access to their acquisition and quantification systems prior to commercialization. Abbelight was not involved in experimental design, data handling, nor in the writing of the manuscript. We also wish to thank the microscopy platform of the Cochin Institute (IMAG’IC), Paris, France for preliminary experiments. We thank S. Dutertre at the Microscopy Rennes Imaging Center (MRic, *Biologie, Santé, Innovation Technologique* -BIOSIT, Rennes, France) for assistance, M. Garfa and all the colleagues at the Cell Imaging Platform, Structure Fédérative de Recherche Necker (Paris, France) for their support. MRic is member of the national infrastructure France-BioImaging supported by the French National Research Agency (ANR-10-INBS-04). This work was supported by the *Centre National de la Recherche Scientifique* (CNRS), an intramural BIOSIT grant, grants from the *Ligue Contre le Cancer Comités d’Ille et Vilaine, des Ĉotes d’Armor et du Finistère*, the *Association pour la Recherche Contre le Cancer* (ARC), and an intramural technology-transfer grant from the *Réseau Technologique de microscopie optique* (RTmfm) to G.B.

## Notes

### Competing Interest Statement

The authors have declared no competing interest.

